# First *De novo* whole genome sequencing and assembly of mutant *Dendrobium* hybrid cultivar ‘Emma White’

**DOI:** 10.1101/2022.06.25.497579

**Authors:** Rubina Sherpa, Ramgopal Devadas, P Suprasanna, Sadashiv Narayan Bolbhat, Tukaram Dayaram Nikam

**Affiliations:** Department of Botany, Annasaheb Awate College, Manchar, Ambegoan 410503, Maharashtra, India; ICAR-National Research Centre on Orchids, Pakyong 737106, Sikkim, India; Nuclear Agriculture and Biotechnology Division, BARC, Mumbai 400085, Maharashtra; Savitribai Phule Pune University, Pune 411007, Maharashtra, India

**Keywords:** Plant science, Plant genetics, Genomics, Dendrobium orchids

## Abstract

*Dendrobium* hybrid cultivar ‘Emma White’ is an ornamental orchid with high commercial demand. We have used gamma-ray induced ‘early flowering mutant’ to generate draft genome sequences with total length (bp) with 678,650,699 and contigs at 447,500 with N50 at 1,423 using the Illumina HiSeqX10 sequencing platform. Here, we report first *de novo* whole genome sequencing and assembly of an early flowering gamma mutant of Emma White hybrid for developing a genomics resource for further studies. The GC content of genome was 33.48%, and predicted 95,529 genes from contig assembly. The predicted genes from the MaSuRCA (version 4.0.3) assembled contigs, when compared with uniprot database using BLASTX program with e-valuecutoff of 10^−3^ resulted 60,741 potential genes governing different pathways in terms for molecular functions, biological process and cellular components. We also identified 216,232 SSRs and 138,856 microsatellite markers. Chromosome level genome assembly of *Dendrobium huoshanense* species was used to RagTag scaffold the available contigs of mutant, where it contained total length of 687,254,899 bp with N50 value 2,096. Largest contiguous length was found with 18,000,059 bp from 30,571 bp. The genome completeness for Emma White RagTag scaffold assembly was assessed to be 93.6% complete using BUSCO v5.2.1 against the Viridiplantae odb10 orthologous dataset. *De novo* whole genome sequencing of gamma mutant Dendrobium hybrid cultivar Emma White (10Gy) isolate was deposited to National Centre for Biotechnology Information (NCBI) with SRA accession <u>SRR16008784</u>, Genebank assembly accession GCA_021234465.1(https://www.ncbi.nlm.nih.gov/assembly/GCA_021234465.1#/st) and Transcriptional Shot Gun assembly accession <u>GJVE00000000</u> under BioProject ID <u>PRJNA763052</u>. This study could provide valuable information for investigating the potential mechanisms of mutation, and guidance for developing Dendrobium hybrid cultivars using mutation breeding.

## Data Description

### Background & Context

The genus *Dendrobium* belongs to theTribe Podochileae, Subtribe Dendrobiinae (Chase et al., 2015). There are about 1,200 species in the genus *Dendrobium*, distributed throughout Southeast Asia and the Southwest Pacific islands. *Dendrobium* has a genome size of 1C = 0.75–5.85 pg (Jones et al., 1998) with diploid chromosome number, 2x=38 (Wang and Xu, 1989). *Dendrobium* hybrids are important flowers of Orchids that has commercial and medicinal demand and potential. Thailand exports 70% of Dendrobiums, with global value of 63.6 billion US$ (Thammasiri, 2015). These Dendrobiums occupy 2^nd^ highest place among sales of potted flowering plants, which accounts 20% contribution from orchid plants in USA (USDA, 2012). Novelty breeding in *Dendrobium* is limited due to narrow genetic makeup of hybrids from *Dendrobium phalaenopsis* (Baker and Baker, 1996) that geographically native to Australia, and inter-sectional cross incompatibility issues to transfer favourable genes (Devadas *et al*., 2016). Reverse genetics through TILLING (Target Induced Local Lesions in Genomics) strategies could offer quick solution to the trait improvement through mutation plant breeding (Jankowicz-Cieslak *et al*., 2017).

*Dendrobium nobile* known as noble orchid and official state flower of Sikkim state (Lucksom, 2007). Complete chloroplast genome of nobile orchid was deciphered recently (Konhar et al., 2016). Biosynthetic pathway of alkaloids for medicinal uses based on functional genomics was extensively studied and reported in *Dendrobium* (Zheng *et al*., 2018; Mou *et al*., 2021). In order to deal with large number of species in *Dendrobium*, large number DNA barcoding systems have been developed and tested aiming for conservation and authentication (Singh et al., 2012; Xu et al., 2015). However, the whole genome sequencing and assembly of *Dendrobium* genus has been reported in four species (Zhang et al., 2021; Zhang et al. 2016) of economic importance for medicinal value only with NCBI, which restricts understanding of phylogenetic diversity among species and their relationship at both inter &intra level to use in crop improvement programmes.

So far, there is no report available for sequence assembly of Dendrobium hybrid cultivars (Taxonomy ID: 136990) and mutants developed (https://www.ncbi.nlm.nih.gov/genome/?term=Dendrobium). In our mutation breeding studies, we have applied Gamma radiation to induce mutations for new variability for orchid genetic improvement. We have chosen a popular and highly adaptable Dendrobium hybrid cultivar ‘Emma White’ derived from complex cross through series of hybridization programme using five Dendrobium species viz., *Dendrobium phalaenopsis* (6 times), *Dendrobium tokai* (1 time), *Dendrobium stratiotes* (1 time), *Dendrobium gouldii* (2 times) and *Dendrobium lineale* (1 times) as parents in pedigree since 1938 to 2006. It was developed by T Orchids, Malaysia registered with Royal Horticultural Society (RHS) in 2006 (http://apps.rhs.org.uk/horticulturaldatabase/orchidregister/orchiddetails.asp?ID=135363). Dendrobium ‘Emma White’ hybrid cultivar a highly cross compatible variety, when used as female parent in hybridization programmes (Devadas et al., 2016). It easily responds *to in-vitro* studies than other hybrids (Devadas et al., 2017), and it was used as one parent in to develop new Dendrobium breeding line, NRCO-42 registered as accession INGR 10073with ICAR-National Bureau of Plant Genetic Resources, India (Devadas et al., 2009). We developed a draft genome sequence of gamma mutant of *Dendrobium* hybrid cultivar ‘Emma White’ (Figure 1), for the first time, which would assist for genetic improvement through deciphering TILLING strategies in future.

**Fig 1:**
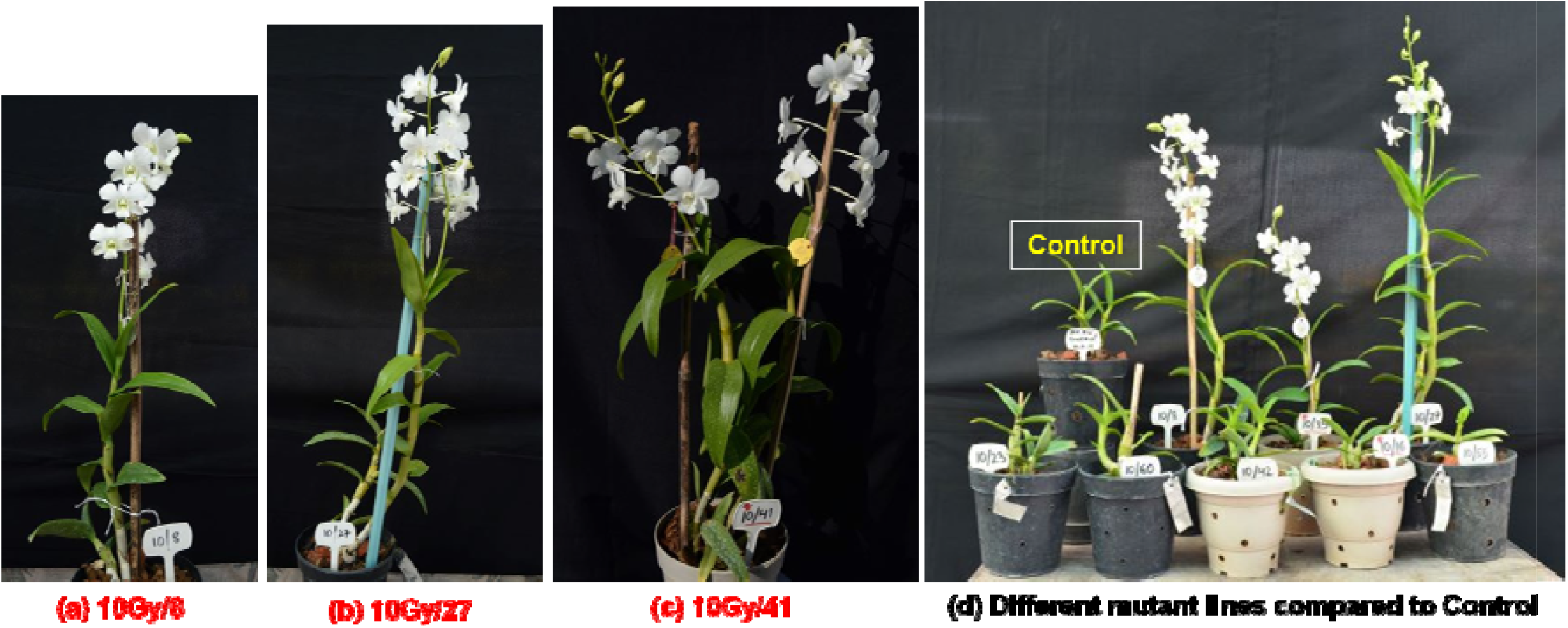
Early flowering mutant lines (10*Gy*) of Dendrobium hybrid cultivar ‘Emma White’ (a) 10Gy/08 (b) 10Gy/27 (c) 10Gy/41 early flowering mutant lines. (d) Comparison of Control plants with flowering mutant plants

**Figure 2:**
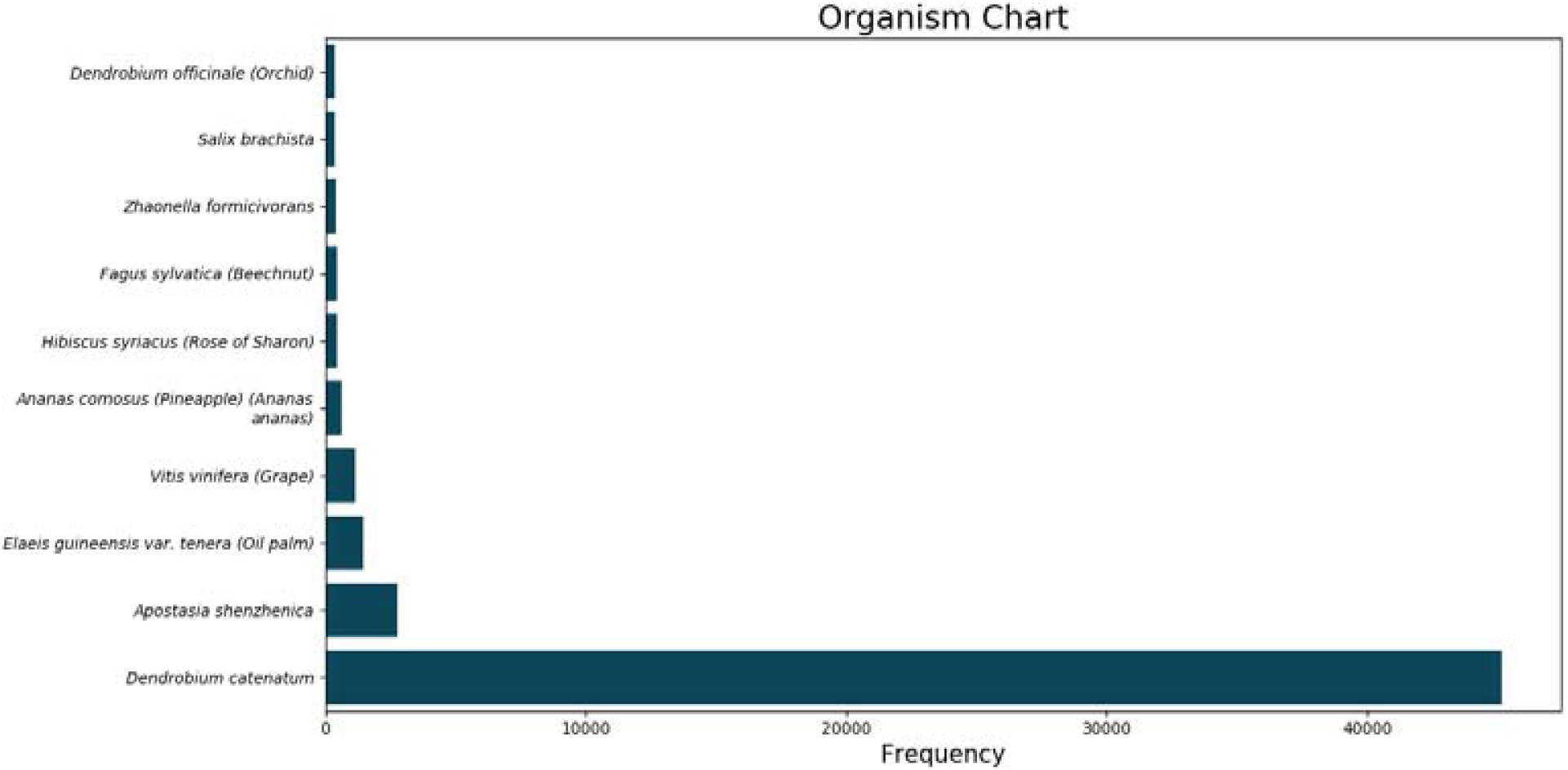
BLASTX hit based on functional predicted gene model for organism

## Methods

### Sampling and DNA preparation

Protocorm Like Bodies (PLBs) of Emma White hybrid were irradiated with gamma @ 10 to 40 *Gy* to induce random mutations@ 32.54 Gy/min using ^60^Cogamma irradiator (Gamma Chamber 5000) at Bhabha Atomic Research Centre, Mumbai as per standard protocols (deAlmeida *et al*., 2014). PLBs were cultured *in-vitro* up to M1V5 generation and plantlets were raised from 10, 20 and 40 *Gy*. Subsequently all the surviving plantlets generated were shifted for hardening and then were grown in polyhouse conditions for phenotypic evaluation. Early flowering mutants were identified among 10 *Gy* plants with several positive traits viz., plant height, pseudo stem length, leaf number, leaf size and spikes during flowering; when there was no flowering (or) delayed flowering was observed in the case of Control, 20 Gy and 40 Gy mutant plants.

Genomic DNA was isolated using the CTAB method (Doyle and Doyle, 1990) from 10 mg of fresh leaves of the first mutant plants during flowering. A DNA sequencing libraries were prepared using a DNA library preparation kit (NEB NextUtra) and tested with Agilent Tapestation for quality validation. DNA fragmentation was performed according to the manufacturer’s instructions to produce fragments having an average length of 150 bp, followed by 5′ and 3′ adaptor ligation. Paired end sequencing was performed using Illumina HiSeqX10 platform. The raw reads were used for *De novo* assembly using assembler MASURCA (version-4.0.3) with default parameters (Zimin *et al*., 2013) with Adapter Removal V2 (Schubert et al., 2016) to get rid of the adapters, low quality reads and bases. Assembly statistics were made using QUAST version 5.2.2 (Gurevich *et al*., 2013) and the levels of conserved genes generated using BUSCO version 5.2.2 (Simao *et al*., 2015). Rag Tag scaffolding was constructed with default parameters running RagTag (v2.1.0) using chromosome based genome assembly of *Dendrobium huoshanense* species available with NCBI (https://www.ncbi.nlm.nih.gov/assembly/GCA_016618105.1). Further, we have performed Benchmarking Universal Single Copy Orthologs (BUSCO) v5.2.2 on the scaffolded fasta with viridiplantae_odb10 lineage dataset to evaluate the assembled scaffold quality and comparison was made (Table 2). Simple sequence repeats of each scaffold were identified by using MISA (v-2.1) script (Thiel *et al*., 2003) and primer designing was done on the predicted SSRs using primer3 (v2.3.6) with default parameters (Rozen and Skaletsky, 2000). MaSuRCA (version 4.0.3) assembled contigs were used for gene prediction model AUGUSTUS (Zimin *et al*., 2017). The predicted genes were compared with uniprot database using BLASTX program with e-value cut off for identification of potential genes governing different pathways. The best BLASTX hit based on query coverage, identity, similarity score and description of each gene was filtered out using custom-made python script and gene ontology was assigned.

## Results &Discussion

We have generated draft genome sequence information by using a gamma-ray radiation induced mutant identified for ‘early flowering’. So far, we have not come across any genome assembly information of whole or partial genome for any modern *Dendrobium* hybrid cultivar (or) mutant *Dendrobium* cultivar using NGS technique. The paired end sequencing using Illumina HiSeqX10 sequencing platform generated 17x genome coverage with 79,792,942 reads (150 bp reads) for 10Gy46 gamma mutant line. NCBI taxonomical analysis of percent alignment on the basis of SRA data matched limited to 8.30% with its closest species *Dendrobium catenatum*, followed by *Phalaenopsis equestris* at 0.55% (NCBI online data for SRA: **SRR16008784**). The genome assembly resulted 678,650,699 bp long, having total 635,396 contigs with longest 30,571 and shortest 300 with mean value 1,068. The N_50_ values 1,423 and GC (%) content 32.48 with contigs 447,500 (Table 1). Rag Tag scaffolding of mutant assembly based on reference genome of *Dendrobium huoshanense* contained total length of 687,254,899 bp increased by 8,604,200 bp with N50 value 2,096 (Table 1). The largest contig length of RagTag scaffolded assembly increased to 18,000,059 from 30,571 with increased N’s per 100 kbp to 1394.82 from ‘0’. The predicted 96,529 genes from the MaSuRCA assembled contigs, when compared with uniprot database using BLASTX program with e-value cut off of 10^−3^ resulted in 60,741 potential genes governing different pathways in terms of Molecular functions, Cellular components and Biological process. BLAST results were filtered based on cut off qcov > 60% and pi identity > 70% to ensure confidence of annotations. We also identified 216,232 SSRs and designed 138,856 microsatellite primers, which can help to assist to generate polymorphic differences among progenies and putative gamma mutant lines. Majority of BLASTX hit has shown affinity to the *D*. *catenatum* species based on functional annotation of genes (Figure 3). BUSCO (version 5.2.2) analysis reveals 913 (56.57%) single-copy orthologs doesn’t match with any data bases indicates the possible impact from both genome back ground of developed hybrid cultivar and also influence of gamma radiation. First, the genome of ‘Emma White’ hybrid cultivar of Dendrobium derived from five unique and unrelated species is complex genome and continuously hybridized repeatedly 11 times over a period of 68 years with selection process for targeted economic trait improvement (Table 2). Low BUSCO values may be related to the fragmented assembly. However, the presence of genome material from several other species of same genus (otherwise contaminant species) in the hybrid cultivar may have resulted drastic changes in the missing BUSCO values (Veeckman et al., 2016; Manni et al., 2021; Waterhouse et al., 2018). Taxonomical analysis of mutant *Dendrobium* based on raw sequence data also supports the view, due to limited synteny with its closest *Dendrobium catenatum* species at below 9%. In addition to these factors, multi-genome hybrid cultivars are genetically heterogeneous with largely outcrossing nature indicating higher compatibility. For example, in case of *Arabidopsis lyrata* an outcrossing species predicted 32,670 genes even at 8.3x DNA coverage, in compared to 27,025 genes in selfing species *Arabidopsis thaliana* (125 Mb) which was separated 10 million years ago (Hu et al., 2011) due loss of genome and rearrangement. In the similar way these novel hybrid cultivars resulted a distinct genome due to introgression from other wild species chosen by Plant Breeders expecting to create new variations in short span of time. Second reason, it can be attributed to deletions, mostly non-coding DNA &transposons and presence of highly mutagenized back ground with severe developmental abnormalities; apart from presence of unclustered genes.

**Table 1:** Genome assembly statistics of gamma mutant of Dendrobium hybrid and RagTag scaffolding.

| Assembly | $\gamma$ mutant<br>(*Dendrobium*<br>hybrid) | *D. catenatum* | *D. huoshanense* | RagTag scaffold<br>of $\gamma$ mutant |
| --- | --- | --- | --- | --- |
|  | (1) | (2) | (3) | RagTag.Scaffold |
| # contigs ($\geq 0$ bp) | 635,396 | 286,396 | **2,256** | 549,354 |
| # contigs ($\geq 1000$ bp) | 213,573 | 29,592 | **2,256** | 163,119 |
| # contigs ($\geq 5000$ bp) | 8,302 | 3,709 | **1,279** | 5,190 |
| # contigs ($\geq 10000$ bp) | 625 | 2,523 | **907** | 370 |
| # contigs ($\geq 25000$ bp) | 7 | 1,684 | **448** | 29 |
| # contigs ($\geq 50000$ bp) | 0 | 1,401 | **145** | 20 |
| Total length ($\geq 0$ bp) | 678,650,699 | 1,104,259,548 | **1,284,285,095** | 687,254,899 |
| Total length ($\geq 1000$ bp) | 439,221,924 | 1,016,149,702 | **1,284,285,095** | 471,383,498 |
| Total length ($\geq 5000$ bp) | 57,139,593 | 973,286,060 | **1,282,134,848** | 184,746,937 |
| Total length ($\geq 10000$ bp) | 7,915,569 | 964,906,549 | **1,279,530,669** | 154,097,582 |
| Total length ($\geq 25000$ bp) | 194,559 | 952,168,543 | **1,272,192,375** | 149,748,116 |
| Total length ($\geq 50000$ bp) | 0 | 942,392,114 | **1,262,665,926** | 149,476,984 |
| # contigs | 447,500 | 64,087 | **2,256** | 369,938 |
| Largest contig | 30,571 | 33,291,853 | **100,197,051** | 18,000,059 |
| Total length | 604,787,319 | 1,040,039,458 | **1,284,285,095** | 616,868,961 |
| GC (%) | 33.48 | 34.61 | **35.73** | 33.49 |
| N50 | 1,423 | 1,149,703 | **71,787,458** | 2,096 |
| N75 | 949 | 434,049 | **52,753,504** | 1,039 |
| L50 | 105,200 | 184 | **8** | 46,924 |
| L75 | 228,327 | 553 | **13** | 154,552 |
| # N's per 100 kbp | 0.00 | 4167.3 | **123.03** | 1394.82 |

(1) https://www.ncbi.nlm.nih.gov/assembly/GCA_021234465.1#/st,
(2) https://www.ncbi.nlm.nih.gov/assembly/GCF_001605985.2
(3) https://www.ncbi.nlm.nih.gov/assembly/GCA_016618105.1

**Table 2:** Overview of BUSCO results of mutant Dendrobium hybrid cultivar and RagTag scaffolded assembly.

| BUSCO Values | $\gamma$ mutant<br>Assembly | | RagTag.Scaffold<br>Assembly | |
| --- | --- | --- | --- | --- |
|  | Number | (%) | Number | (%) |
| Complete BUSCOs (C) | 286 | 16.60 | 312 | 73.4 |
| Complete and single-copy BUSCOs (S) | 249 | 15.43 | 306 | 72.0 |
| Complete and duplicated BUSCOs (D) | 19 | 1.18 | 6 | 1.4 |
| Fragmented BUSCOs (F) | 433 | 26.83 | 86 | 20.2 |
| Missing BUSCOs (M) | 913 | 56.57 | 27 | 6.4 |
| Total BUSCO groups searched | 1,614 | 100 | 425 | 100 |

The mutant Dendrobium hybrid sequencing and genome assembly can be adopted as primary reference genome and complement existing conventional Dendrobium species in public domain. These studies on induced mutants provide discovery of new alleles rapidly at low cost using high-throughput tilling (Kurowska et al., 2011), especially in vegetatively propagated Dendrobium hybrids, for obtaining high density mutations using gamma mutation breeding as evident from other crops (Datta et al., 2018; Suprasanna et al., 2015). The results provide a baseline for further research on the molecular understanding of desired traits in the mutant germplasm and to develop genome resources for use in orchid improvement.

## Data availability

*De novo* whole genome sequencing of gamma mutant Dendrobium hybrid cultivar Emma White (10Gy) isolate was deposited to National Centre for Biotechnology Information (NCBI) with SRA accession <u>SRR16008784</u> and Gene bank assembly accession GCA_021234465.1(https://www.ncbi.nlm.nih.gov/assembly/GCA_021234465.1#/st) and and Transcriptional Shot Gun assembly accession <u>GJVE00000000</u> under BioProject ID <u>PRJNA763052</u> available in the public domain with access.

## Ethical approval

Not applicable

## Consent for publication

Not applicable

## Competing interests

No competing interests were disclosed.

## Grant information

The authors(s) declare that no special grants were involved specific to this mutant sequencing. This work has been carried by Scholar for validation of research findings.

## Author’s contribution

RS conducted the gamma radiation experiment, mutant plant development and genome analysis. RD prepared concept and research formulation of project and wrote the manuscript. PS assisted radiation facilities and re-edit manuscript. SNB and TKN assisted the overall supervision of work and interpreted the data. All authors read, edited and approved the manuscript submission.

## Acknowledgements

RS and RD are thankful to NABTD, BARC, Government of India for providing radiation facilities. RD is thankful to BRNS, Government of India for proving fellowship to scholar and funding to core project. The authors are thankful to Director, ICAR-NRC on Orchids for providing institutional support. M/s AgriGenome Labs Private Limited, Kerala, India for sequencing work.

## Notes

### Competing Interest Statement

The authors have declared no competing interest.

